# Substrate trapping approach identifies TRIM25 ubiquitination targets involved in diverse cellular and antiviral processes

**DOI:** 10.1101/2022.03.17.484727

**Authors:** Emily Yang, Serina Huang, Yasaman Jami-Alahmadi, Gerald M McInerney, James A Wohlschlegel, Melody MH Li

## Abstract

The tripartite motif (TRIM) family of E3 ubiquitin ligases is well known for its roles in antiviral restriction and innate immunity regulation, in addition to many other cellular pathways. In particular, TRIM25-mediated ubiquitination affects both carcinogenesis and antiviral response. While individual substrates have been identified for TRIM25, it remains unclear how it regulates diverse processes. Here we characterized a mutation, R54P, critical for TRIM25 catalytic activity, which we successfully utilized to “trap” substrates. We demonstrated that TRIM25 targets proteins implicated in stress granule formation (G3BP1/2), nonsense-mediated mRNA decay (UPF1), and nucleoside synthesis (NME1). R54P abolishes TRIM25 inhibition of alphaviruses independently of the host interferon response, suggesting that this antiviral effect is a direct consequence of ubiquitination. Consistent with that, we observed diminished antiviral activity upon knockdown of several TRIM25-R54P specific interactors including NME1. Our findings highlight that multiple substrates mediate the cellular and antiviral activities of TRIM25, illustrating the multi-faceted role of this ubiquitination network in diverse biological processes.

## INTRODUCTION

Addition of ubiquitin, or ubiquitination, is a post-translational modification that is highly conserved in eukaryotic organisms, and operates in myriad cellular pathways. Ubiquitin is a small, 76 amino acid protein that must be activated by E1 enzymes, passed to E2 carrier enzymes, and finally covalently attached to lysines on substrates by E3 ligases. Though only one enzyme is needed at each step, their numbers vary widely. Humans encode 2 E1 enzymes, about 40 E2 enzymes, and upwards of 600 E3 ligases.^1,2^ This vast number of E3 ligases is needed because they determine substrate specificity; however, the means by which E3 ligases identify their substrates and the array of substrates ubiquitinated by any given E3 ligase remain largely unknown.

The tripartite motif containing protein (TRIM) family is one of the largest families of E3 ligases, with over 70 *TRIM* genes in humans.^3^ TRIMs share three common domains at their N-terminus – the catalytic RING domain, 1 to 2 B-Box domains, and a coiled-coil domain – but differ in their C-termini.^3^ These varied C-termini determine TRIM substrate specificity, allowing this large family of proteins to regulate diverse cellular processes, including but not limited to viral restriction, immune signaling, stress responses, proliferation, and differentiation.^4–7^ Mutations in *TRIM* genes have been associated with rare genetic diseases, including developmental, muscular, and neurological disorders.^8,9^ However, development of targeted therapeutic approaches has been hindered by not only the lack of knowledge on their specific substrates, but also the frequent involvement of TRIMs in multiple cellular processes. One prime example is TRIM25, which functions in both cancer and antiviral innate immunity.^10,11^ When examined in the context of cancer, TRIM25-mediated ubiquitination primarily targets varied proteins for proteolytic degradation, which can either enhance or hinder carcinogenesis.^12–16^

Many of the TRIM proteins are upregulated by interferon (IFN) and play significant roles in the host innate immune response.^7^ Upon detection of viral infection by the host cell, type I IFN is produced, inducing expression of hundreds of IFN-stimulated genes (ISGs) to establish an antiviral environment.^17,18^ TRIM25 is one such ISG which not only stimulates innate immune signaling by ubiquitinating and activating the dsRNA sensor RIG-I, but also functions as a critical co-factor of another ISG, zinc finger antiviral protein (ZAP).^19–21^ While TRIM25 has been shown to complex with ZAP in the context of several different viral infections,^22^ its ligase activity has only been tied to its participation in blocking translation of incoming RNA genomes of alphavirus (family *Togaviridae*).^20^ Given that ubiquitination of ZAP or lack thereof fails to affect its viral translation inhibition,^20^ it is likely that TRIM25 antiviral involvement depends on its ubiquitination of other cellular proteins. Interestingly, both TRIM25 and ZAP not only bind viral RNA but also interact with other RNA binding proteins, implying that proteins involved in RNA processes may feature prominently among TRIM25 substrates.^23–26^

In light of this question, we set out to identify novel TRIM25 substrates that may play a role in translation and RNA processes. Because identification of E3 ligase substrates is technically challenging due to the transient nature of ligase-substrate interactions, we utilized a “substrate trapping” approach as previously reported^27^ to capture TRIM25 interactors in a co-immunoprecipitation (IP)/mass spectrometry (MS) experiment. We sought to generate a TRIM25 mutant that would be unable to interact with the upstream E2 carrier enzyme, thus simultaneously rendering it incapable of ubiquitination and prolonging its interactions with substrates. We identified a point mutation, R54P, in the TRIM25 RING catalytic domain, which almost completely abolishes its autoubiquitination in cells.

We found that TRIM25-R54P enriches for additional interactors as compared to TRIM25-WT, though almost all of the more highly enriched interactors are shared by both TRIM25-WT and -R54P. Further characterization of some of the most highly enriched interactors, Ras-GTPase-activating protein SH3-domain binding proteins (G3BP) 1 and 2, RNA helicase up-frameshift protein 1 (UPF1), and nucleoside diphosphate kinase 1 (NME1), has validated their identification as novel TRIM25 substrates. Moreover, upon characterization of its antiviral activity, the TRIM25-R54P mutant demonstrates a complete loss of inhibition against a panel of Old World and New World alphaviruses albeit higher IFN and ISG expression compared to WT, suggesting that ubiquitination of TRIM25 substrates directly leads to activation of an antiviral state. Altogether, we have identified both known and novel interactors as TRIM25 substrates, and demonstrated the validity of this “substrate trapping” approach in identifying bona fide E3 ligase substrates. We have shed light on the ways that TRIM25-mediated ubiquitination might target substrates to modulate translation, nucleic acid metabolism, and antiviral response, paving the way for further work characterizing the critical role of TRIMs in diverse cellular and viral processes.

## RESULTS

### Point mutations in TRIM25 RING domain almost completely abolish TRIM25 autoubiquitination

It is technically challenging to identify E3 ligase-substrate interactions as they are often transient, resulting in proteasomal degradation or a change in localization or activity of the substrates. In order to enrich for transient E3 ligase-substrate interactions, we turned to a less conventional co-IP approach that makes use of E3 mutants unable to interact with E2 conjugating enzymes. This prevents ubiquitin transfer to E3 substrates and their subsequent targeting to other cellular pathways and as a result, “trapping” these substrates. This approach successfully identified the cellular ‘structural maintenance of chromosomes’ (Smc) complex Smc5/6 as being targeted by hepatitis B virus X protein for ligase-mediated degradation.^27^ We hypothesized that a similar approach would serve to identify TRIM25 substrates, which will be immunoprecipitated more robustly with a TRIM25 E2 binding mutant than with TRIM25-WT, as the former is unable to mediate transfer of ubiquitin from E2 to substrates.

Residues important for the RING-E2 interaction and thus necessary for ligase activity have already been identified in the RING E3 ligase MDM2.^28^ We aligned the structure of the TRIM25 RING domain complexed to E2-ubiquitin (Ub) to the analogous MDM2-E2-Ub structure and identified two conserved critical E2 interaction residues in TRIM25 RING, I15 and R54 (Fig. 1A). To assess loss of ligase activity, we transfected HA-tagged Ub and FLAG-tagged TRIM25 into 293T cells and immunoprecipitated TRIM25 in denaturing conditions. We then blotted for HA-Ub, wherein polyubiquitination manifests as a ladder of bands. These TRIM25 E2 binding mutants (I15K and R54P), are deficient in autoubiquitination, suggesting successful crippling of ligase activity (Fig. 1B). Individual E2 binding mutants retain a mono-Ub band (Fig. 1B), so we generated the double mutant I15K/R54P, which did not display further reduction in ligase activity (Fig. 1B). Therefore, we selected the R54P mutant for future co-IP/MS studies since this mutation has previously been shown to reduce TRIM25 catalytic activity and polyubiquitin chain formation.^29^

**Figure 1.**
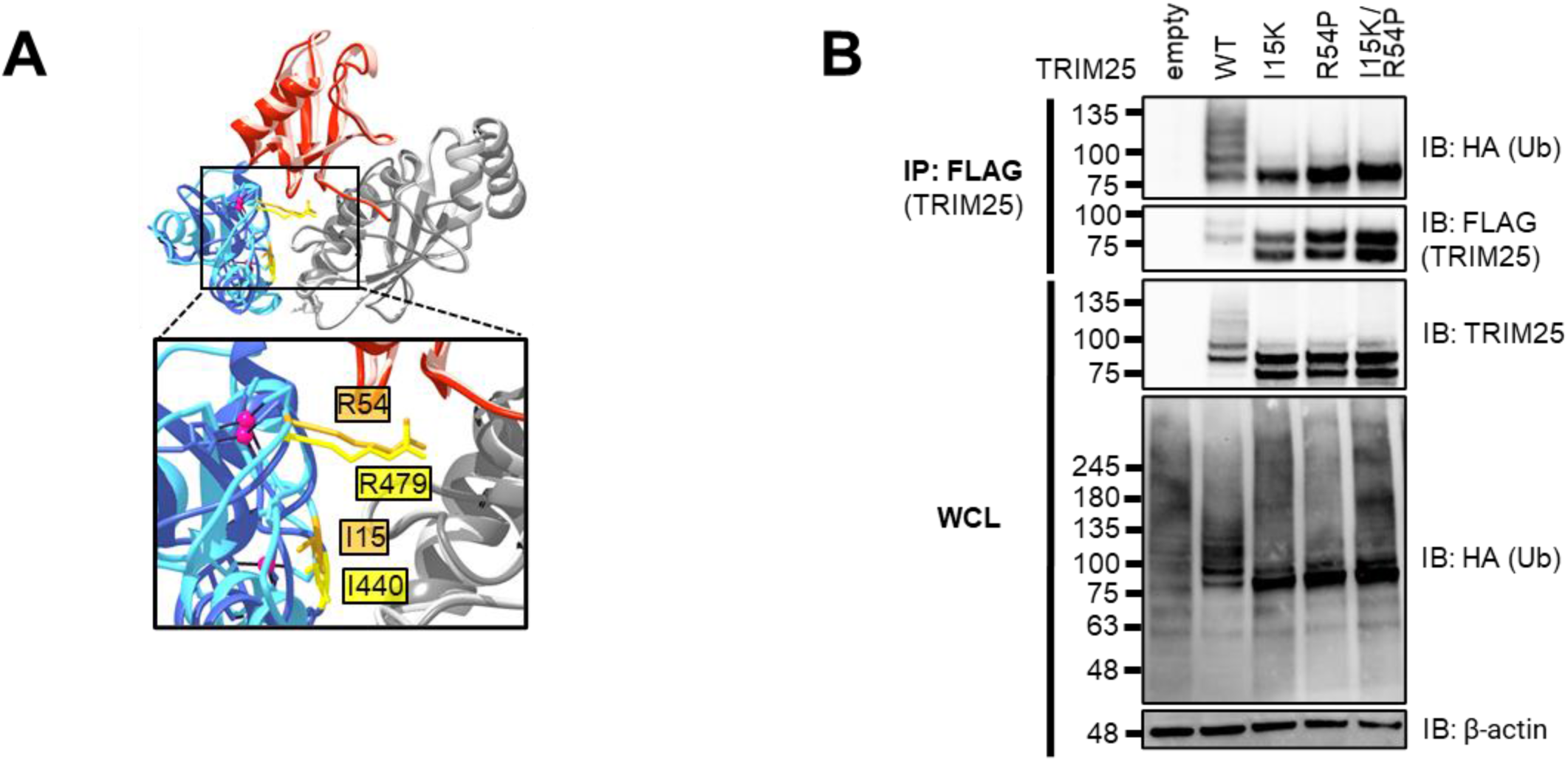
Individual TRIM25 RING residues required for TRIM25 autoubiquitination. **(A)** Alignment of the RING E3 ligases MDM2 (dark blue) and TRIM25 (light blue) in complex with ubiquitin (red) and the E2 UbcH5 (gray), performed using UCSF Chimera.^82^ Highlighted in gold (TRIM25) and yellow (MDM2) are homologous residues. PDB: 5MNJ (MDM2), 5EYA (TRIM25). **(B)** Western blot of lysates of 293T cells transfected with FLAG-TRIM25 mutants and HA-ubiquitin (Ub). Data representative of three independent experiments.

### Substrate trapping approach enriches for novel TRIM25 interactors

Next, we asked what proteins are modified by TRIM25, as identification of these substrates will elucidate how ubiquitination facilitates TRIM25-mediated cellular and antiviral activities. We first used CRISPR-Cas9 to generate a TRIM25 KO 293T cell line (Fig. S1). We then stably integrated doxycycline (dox) inducible FLAG-tagged TRIM25 wild-type (WT) and mutant R54P using the ePiggyBac (ePB) transposon system,^30^ where both TRIM25-WT and TRIM25-R54P are similarly induced in a dose-dependent manner (Fig. 2A). TRIM25 protein levels are comparable upon detection using a FLAG or TRIM25-specific antibody (Fig. 2A).

**Figure 2.**
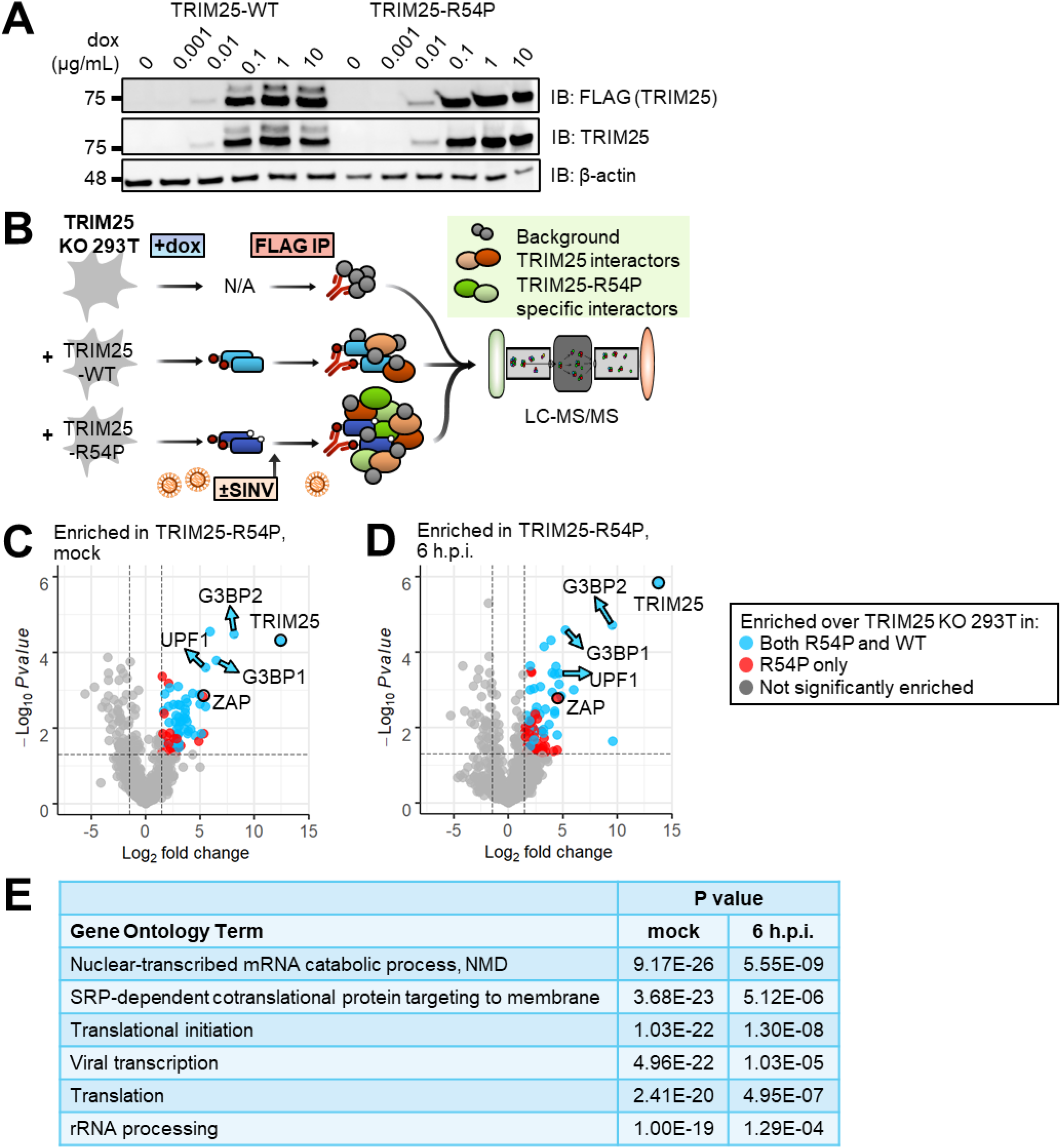
TRIM25 co-IP/MS Identifies TRIM25 interactors. **(A)** Western blot of TRIM25 inducible 293T cell lines in the presence of increasing amount of dox (0, 0.001, 0.01, 0.1, 1, and 10 μg/mL). Data are representative of two independent experiments. **(B)** Schematic of co-IP/MS experiment to identify TRIM25 interactors. **(C-D)** Volcano plots of proteins significantly enriched over TRIM25 KO background in TRIM25-R54P co-IP/MS in the **(C)** absence or **(D)** presence of viral infection. Data representative of two independent experiments. Blue dots represent proteins that were also significantly enriched in TRIM25-WT co-IP and red dots represent proteins that were only enriched in TRIM25-R54P co-IP. Proteins were counted as enriched when log2FC>1.5 and - log10Pvalue>1.3 (Pvalue<0.05). The R package EnhancedVolcano^83^ was used to generate volcano plots. **(E)** Gene ontology terms significantly enriched in all unique TRIM25-WT and TRIM25-R54P interactors. Analysis performed for GO terms in biological processes using DAVID.^31,32^

To capture TRIM25 substrates, we performed two independent co-IP/MS experiments using our reconstituted TRIM25 KO 293T cell lines (Fig. 2B). We induced TRIM25-WT or -R54P expression in the presence or absence of the prototype alphavirus Sindbis virus (SINV), performed a FLAG IP to enrich for TRIM25, and analyzed the resultant protein mixture using MS. TRIM25 KO 293T cells were used as a control. We found that this “substrate trapping” approach enriches for interactors specific to TRIM25-R54P under both mock and infected conditions (Fig. 2C-D, red circles). These TRIM25-R54P-specific interactors tend to have lower fold change in abundance over background than interactors common to both TRIM25-WT and TRIM25-R54P (Fig. 2C-D, blue circles), suggesting that the TRIM25-R54P co-IP/MS captures weaker interactions not identified with TRIM25-WT. After filtering for interactors enriched in both independent experiments, we found that TRIM25-R54P enriches for 14 unique interactors under mock conditions (Table 1) and that almost all TRIM25-WT interactors (25 of 30) are also present as TRIM25-R54P interactors (Table 2), indicating that TRIM25-R54P is otherwise functionally similar to TRIM25-WT. During viral infection, TRIM25-R54P enriches for all TRIM25-WT interactors in addition to 16 unique interactors (Table 3-4), suggesting an effective “substrate trap.” Interestingly, we found that the number of TRIM25 interactors drastically decreases during viral infection for both TRIM25-WT (29 to 7 interactors; Tables 2 and 4) and TRIM25-R54P (38 to 23 interactors; Tables 1 and 3). We used DAVID bioinformatics resources^31,32^ to find that TRIM25 interactors are highly enriched in GO terms involved in translation, RNA metabolism, and viral transcription (Fig. 2E). This is in line with our hypothesis that TRIM25 substrates mediate diverse cellular and viral processes as a consequence of ubiquitination.

**Table 1.** TRIM25-R54P interactors in the absence of virus. Interactors pulled down in both independent experiments shown here; proteins also enriched in TRIM25-WT co-IP are italicized and bolded. Fold change = FC. In EXP #2, the “i” prefacing log2FC and Pvalue refers to how missing data values were imputed.

| Protein | EXP #1<br>log2FC | EXP #2<br>i log2FC | EXP #1<br>Pvalue | EXP #2<br>-log10iPvalue |
| --- | --- | --- | --- | --- |
| **TRIM25** | **12.3906422** | **7.51146541** | **5.11E-05** | **2.12064733** |
| G3BP2 | 8.10501888 | 5.54740584 | 3.34E-05 | 1.50058344 |
| **G3BP1** | **6.50319403** | **4.03438911** | **0.00016672** | **1.7320921** |
| **PABPC1** | **5.9317735** | **5.10019115** | **2.80E-05** | **3.28427084** |
| **UPF1** | **5.49937981** | **4.78953669** | **0.0002495** | **1.43787018** |
| RPL27 | 5.34447755 | 3.69353218 | 0.01445622 | 1.55332697 |
| **ZC3HAV1** | **5.31980842** | **5.31119453** | **0.00149558** | **1.41536917** |
| **ZCCHC3** | **5.12257102** | **3.75470266** | **0.01424379** | **1.7776868** |
| RPL36 | 4.97631769 | 3.47534848 | 0.00130952 | 1.79262008 |
| MOV10 | 4.91240181 | 4.49394342 | 0.02289689 | 1.7977924 |
| **RPLP2** | **4.1266814** | **5.86061559** | **0.01106949** | **1.55003589** |
| **SSB** | **3.89157859** | **4.47911201** | **0.0123085** | **1.70757999** |
| **RPS3A** | **3.78778898** | **2.6935167** | **0.00693553** | **1.89210098** |
| MRPL11 | 3.76533373 | 5.47257077 | 0.00216534 | 1.39579663 |
| RPL3 | 3.65992515 | 2.14601333 | 0.00570166 | 1.79197282 |
| **RPS12** | **3.65715187** | **3.49122989** | **0.00172346** | **1.89210098** |
| **NME1** | **3.61188084** | **4.79415355** | **0.00405771** | **1.79197282** |
| **IGF2BP3** | **3.56496076** | **3.28060717** | **0.01055553** | **1.50058344** |
| **RPS8** | **3.2641646** | **3.27360754** | **0.00648829** | **1.75046739** |
| DNAJA1 | 3.17029213 | 3.92452385 | 0.00248228 | 1.91245992 |
| RPL21 | 3.05781202 | 2.39189309 | 0.03343166 | 1.4733235 |
| **DDX21** | **3.05473047** | **1.88642662** | **0.02565655** | **1.7320921** |
| **RPL7A** | **3.03971967** | **2.5230631** | **0.00882745** | **1.7320921** |
| **DDX50** | **3.03354054** | **3.65683219** | **0.03340781** | **1.52755659** |
| **RPL14** | **3.01017649** | **2.78807831** | **0.01479679** | **1.79197282** |
| **HSPA9** | **2.97612651** | **2.5383849** | **0.0007997** | **1.7320921** |
| **RPL8** | **2.95324948** | **2.63561626** | **0.00240818** | **1.50058344** |
| **RPL4** | **2.94771854** | **1.92716275** | **0.01579881** | **1.49666772** |
| **RPL30** | **2.92460761** | **4.9910042** | **0.01395897** | **1.81397601** |
| RPL6 | 2.87523998 | 1.95453062 | 0.01081102 | 1.49666772 |
| RPL19 | 2.77420542 | 5.26953341 | 0.0455276 | 1.35734257 |
| CXorf56 | 2.61987631 | 2.58742073 | 0.01931576 | 1.34462142 |
| **MRPS25** | **2.55900639** | **3.33809393** | **0.00246282** | **1.54084885** |
| POLDIP2 | 2.24491261 | 2.4533861 | 0.00085532 | 1.64450634 |
| **IGF2BP2** | **2.19855163** | **3.2324213** | **0.00701643** | **1.45704269** |
| **HSPA5** | **2.09668197** | **2.60999743** | **0.00273418** | **1.79262008** |
| NCL | 1.94667851 | 2.58059341 | 0.01659916 | 2.629704 |
| **HSPA8** | **1.63949288** | **2.00543879** | **0.0035694** | **1.79262008** |
| RTRAF | 1.60321627 | 1.76389708 | 0.01444176 | 1.31238098 |

**Table 2.** TRIM25-WT interactors in the absence of virus. Interactors pulled down in both independent experiments shown here; proteins also enriched in TRIM25-R54P co-IP in both independent experiments are italicized and bolded. Fold change = FC. In EXP #2, the “i” prefacing log2FC and Pvalue refers to how missing data values were imputed.

| Protein | EXP #1<br>log2FC | EXP #2<br>ilog2FC | EXP #1<br>Pvalue | EXP #2<br>-log10iPvalue |
| --- | --- | --- | --- | --- |
| **TRIM25** | **12.3454** | **7.211613** | **5.18E-05** | **2.355901** |
| **NME1** | **5.894505** | **5.692311** | **0.000639** | **2.112497** |
| **PABPC1** | **5.178075** | **4.356066** | **4.80E-05** | **2.648698** |
| **UPF1** | **4.154268** | **4.989455** | **0.000742** | **1.561182** |
| **G3BP1** | **3.839632** | **4.375498** | **0.001287** | **2.076582** |
| **SSB** | **3.671733** | **5.281826** | **0.014983** | **1.665287** |
| **RPS12** | **3.521782** | **3.017109** | **0.001987** | **2.238969** |
| **DDX21** | **3.515317** | **1.604128** | **0.01627** | **2.073546** |
| **ZCCHC3** | **3.406602** | **2.467616** | **0.031369** | **1.500504** |
| **MRPL11** | **3.112947** | **4.498654** | **0.004378** | **1.829982** |
| **MRPS25** | **3.112087** | **5.368709** | **0.001176** | **1.805081** |
| **RPS3A** | **3.01824** | **1.917857** | **0.015203** | **2.112497** |
| **RPL14** | **2.90745** | **2.557471** | **0.016614** | **1.710395** |
| **RPS8** | **2.859517** | **3.070731** | **0.010332** | **2.112497** |
| RPL23A | 2.829982 | 2.504147 | 0.03774 | 2.289792 |
| LARP1 | 2.82004 | 2.717784 | 0.017809 | 1.648418 |
| **HSPA9** | **2.78228** | **2.547643** | **0.001036** | **1.977132** |
| **DDX50** | **2.775815** | **5.515807** | **0.041851** | **1.86816** |
| **RPLP2** | **2.7105** | **5.808829** | **0.042541** | **1.359565** |
| **RPL7A** | **2.680142** | **2.051832** | **0.013614** | **2.112497** |
| **RPL8** | **2.671594** | **1.909501** | **0.003493** | **1.458887** |
| **IGF2BP3** | **2.665042** | **3.649089** | **0.027568** | **1.663079** |
| **HSPA5** | **2.358487** | **2.765141** | **0.001759** | **2.05426** |
| FMR1 | 2.163144 | 2.141417 | 0.04163 | 1.562371 |
| **RPL4** | **2.089384** | **1.963929** | **0.046256** | **2.076582** |
| **RPL30** | **1.982778** | **4.991994** | **0.047276** | **2.663368** |
| RPL32 | 1.910939 | 2.096376 | 0.03637 | 2.05426 |
| **IGF2BP2** | **1.731804** | **2.157021** | **0.015972** | **1.449724** |
| **HSPA8** | **1.655745** | **2.262534** | **0.003442** | **1.648418** |
| FXR1 | 1.522065 | 4.862743 | 0.002183 | 1.592811 |

**Table 3.** TRIM25-R54P interactors during viral infection. Interactors pulled down in both independent experiments shown here; proteins also enriched in TRIM25-WT co-IP in both independent experiments are italicized and bolded. Fold change = FC. In EXP #2, the “i” prefacing log2FC and Pvalue refers to how missing data values were imputed.

| Protein | EXP #1<br>log2FC | EXP #2<br>ilog2FC | EXP #1<br>Pvalue | EXP #2<br>-log10iPvalue |
| --- | --- | --- | --- | --- |
| **TRIM25** | **13.8222763** | **6.87172529** | **1.48E-06** | **1.2781953** |
| G3BP2 | 9.49865601 | 7.65732671 | 1.95E-05 | 1.39507974 |
| G3BP1 | 5.24931968 | 2.50691034 | 2.61E-05 | 1.91751582 |
| NME1 | 4.81029586 | 3.42772379 | 0.00145919 | 1.39507974 |
| **H1FX** | **4.64670773** | **4.49007695** | **0.00072161** | **1.46059266** |
| **UPF1** | **4.64084486** | **5.24120822** | **0.0003629** | **1.93751203** |
| ZC3HAV1 | 4.47005094 | 2.7706705 | 0.0016896 | 1.41791083 |
| RPL8 | 4.25732657 | 2.98030126 | 0.00436065 | 1.80121376 |
| PABPC4 | 4.25391021 | 2.67042471 | 0.00037935 | 1.41791083 |
| PABPC1 | 4.24158137 | 3.53536817 | 0.00365513 | 1.44245201 |
| **HP1BP3** | **3.99809257** | **4.08820333** | **0.00036134** | **1.50049849** |
| **LARP1** | **3.92792615** | **4.59386762** | **4.74E-05** | **1.60943756** |
| **DDX50** | **3.73502582** | **4.13736412** | **0.00829299** | **1.47454054** |
| RPL21 | 3.39655083 | 3.45363607 | 0.03185577 | 1.44245201 |
| RPL29 | 3.38014514 | 2.52995575 | 0.0032809 | 2.06147369 |
| **HSPA9** | **3.27541907** | **2.03709492** | **7.07E-05** | **2.0451482** |
| **ZCCHC3** | **3.23306468** | **4.48645336** | **0.01122933** | **1.54364156** |
| RPS3A | 3.18225495 | 1.52708792 | 0.04409253 | 1.37667477 |
| GLYR1 | 3.0147273 | 2.59333015 | 0.03723913 | 1.58508512 |
| YBX1 | 2.60522001 | 1.72691896 | 0.03659547 | 1.44245201 |
| GNL3L | 2.47305963 | 5.27949599 | 0.0044276 | 1.57942545 |
| RPL19 | 2.44163445 | 2.90396027 | 0.0040375 | 1.75087336 |
| MRPS26 | 1.98052486 | 1.90420543 | 0.03219554 | 1.30553169 |
| MRPS9 | 1.58301349 | 2.55803142 | 0.00963025 | 1.37679621 |

**Table 4.** TRIM25-WT interactors during viral infection. Interactors pulled down in both independent experiments shown here; proteins also enriched in TRIM25-R54P co-IP in both independent experiments are italicized and bolded. Fold change = FC. In EXP #2, the “i” prefacing log2FC and Pvalue refers to how missing data values were imputed.

| Protein | EXP #1<br>log2FC | EXP #2<br>ilog2FC | EXP #1<br>Pvalue | EXP #2<br>-log10iPvalue |
| --- | --- | --- | --- | --- |
| **TRIM25** | **13.1932** | **6.428015** | **1.78E-06** | **1.15094** |
| **UPF1** | **3.86158** | **5.511565** | **0.00074** | **1.624772** |
| **DDX50** | **2.443466** | **5.210275** | **0.033602** | **1.456255** |
| **ZCCHC3** | **2.653732** | **4.900796** | **0.016533** | **1.406113** |
| **LARP1** | **2.61938** | **4.613364** | **0.000234** | **1.420394** |
| **HP1BP3** | **2.749535** | **3.838037** | **0.001527** | **1.751598** |
| **H1FX** | **3.948205** | **3.229771** | **0.001347** | **1.744913** |
| **HSPA9** | **2.62801** | **2.3519** | **0.000169** | **1.408544** |

### TRIM25 interacts with G3BP1 and 2 through a conserved binding motif and modifies them with predominantly K63 polyubiquitin chains

Among the most enriched TRIM25-R54P interactors in the presence and/or absence of SINV infection (Tables 1, 3), we identified the core stress granule proteins G3BP1 and 2, RNA helicase UPF1 (Fig. 2C-D, blue arrows), and the metastatic suppressor and nucleoside kinase NME1 as high priority candidates given our interest in RNA metabolic and translation processes (G3BP1 and 2, UPF1) and TRIM25’s role in regulating carcinogenesis (NME1). Next, we asked whether any of these are TRIM25 ubiquitination substrates.

Both G3BP1 and G3BP2, hereafter collectively referred to as G3BP, associate very strongly with TRIM25 in the co-IP/MS (Tables 1-4, G3BP1: log2FoldChange 2.5 – 6.5; G3BP2: log2FoldChange 5.5 – 9.5). G3BP normally function in stress granule (SG) assembly, interacting with RNA and other cellular proteins to induce SG formation.^33,34^ Interestingly, the Old World alphaviruses exploit G3BP to promote their own replication.^35–38^ These viruses utilize their non-structural protein nsP3 to recruit G3BP into viral replication complexes, which disrupts antiviral SG formation,^39^ clusters viral replication complexes,^38^ and recruits translation initiation machinery.^40^ By doing so, alphaviruses enhance viral replication at the cost of endogenous G3BP function.

Previous work identified an FGDF peptide motif in alphavirus nsP3 which binds with high affinity to G3BP.^39,41^ More recent work characterizing viral-host interaction motifs has uncovered a conserved G3BP-binding motif, ΦxFG (where Φ is a hydrophobic residue).^42^ This G3BP interaction motif is present in both viral and host proteins, such as the cellular SG protein and known G3BP interactor USP10, and is remarkably similar to the alphavirus nsP3-G3BP interaction motif, FGDF, but likely binds with lower affinity.^42^ Moreover, TRIM25 was identified as a G3BP1 interaction partner.^42^ Mutating the latter two amino acids in the TRIM25-specific motif (404-PTFG-407), to alanine (404-PTAA-407) was sufficient to abolish TRIM25-G3BP1 interaction.^42^ Meanwhile, TRIM25 and G3BP2 have previously been shown to interact in the context of prostate cancer.^43^ To examine whether this motif is also necessary for TRIM25-G3BP2 interaction, we co-transfected myc-tagged G3BP into TRIM25 KO 293T cells along with FLAG-tagged TRIM25-WT, -R54P, or -PTAA, and performed a FLAG IP to pull down TRIM25. While both TRIM25-WT and -R54P robustly associate with both G3BP1 and −2, TRIM25-PTAA does not associate with either G3BP1 or 2 (Fig. 3A), validating our co-IP/MS identification of G3BP as TRIM25 interactors.

**Figure 3.**
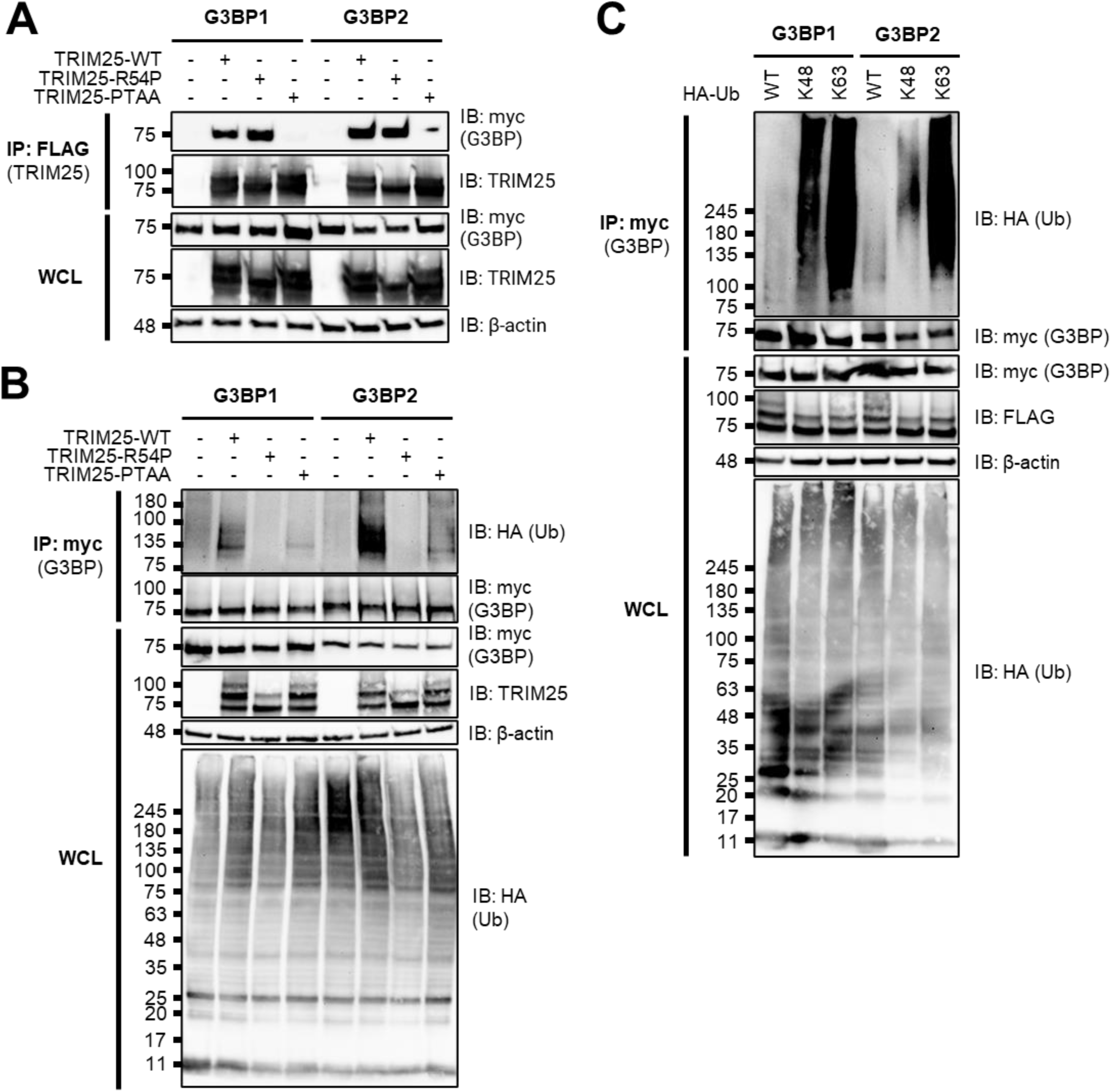
TRIM25 interacts with and ubiquitinates G3BP. **(A)** Western blot of TRIM25 KO 293T cells transfected with myc-G3BP1/2 and FLAG-TRIM25-WT, -R54P, or -PTAA. Lysates were subjected to FLAG IP. Data are representative of three independent experiments. **(B-C)** Western blot of TRIM25 inducible cells transfected with myc-G3BP1/2 and HA-Ub-WT, or **(C)** −K48, or −K63. Lysates were subjected to myc IP. Data are representative of three independent experiments.

We then used the ePiggyBac transposon system to reconstitute TRIM25 KO 293T cells with dox inducible TRIM25-PTAA. To establish that TRIM25 ubiquitinates G3BP and that the TRIM25-G3BP interaction is necessary for ubiquitination, we co-transfected myc-tagged G3BP with HA-Ub into TRIM25-WT, -R54P, and -PTAA inducible cell lines. After inducing TRIM25 expression, we performed a myc IP and probed for the presence of ubiquitinated G3BP. We found that both G3BP1 and 2 are robustly polyubiquitinated only in the presence of TRIM25-WT (Fig. 3B), again validating our co-IP/MS identification of G3BP as TRIM25 substrates. No ubiquitination is detected in the presence of ligase-deficient TRIM25-R54P, whereas ubiquitination is dramatically diminished in the presence of G3BP-interaction deficient TRIM25-PTAA (Fig. 3B). Interestingly, TRIM25 appears to more robustly ubiquitinate G3BP2 as compared to G3BP1 (Fig. 3B). Given the TRIM25-mediated polyubiquitination of G3BP1 and 2, we then characterized G3BP ubiquitination linkage type. To do so, we transfected our TRIM25-WT inducible cell line with myc-G3BP1 or −2 and different forms of HA-Ub; −WT, - K48, and −K63. Ub-K48 and −K63 have all lysines mutated to arginine except −K48 and −K63, respectively, such that only K48 or K63 polyubiquitin chains are able to be formed.^44^ We found that both G3BP1 and 2 are most robustly ubiquitinated in the presence of Ub-K63, suggesting that TRIM25 primarily mediates K63-linked ubiquitination of both proteins (Fig. 3C). Interestingly, while both G3BP1 and 2 exhibit a lower level of ubiquitination in the presence of Ub-K48, G3BP1 possesses more K48-linked polyubiquitin chains as compared to G3BP2 (Fig. 3C), indicating that TRIM25 is able to distinguish between and differentially ubiquitinate these related proteins.

### TRIM25 interacts with and mono-ubiquitinates UPF1 at K592

Moreover, UPF1 associates very strongly with TRIM25 in the co-IP/MS (Tables 1-4, log2FoldChange 3.9 – 5.5), supporting a role for UPF1 as a novel TRIM25 interactor. UPF1 is best known for its central role in nonsense-mediated mRNA decay (NMD), where it is recruited to premature termination codons to catalyze the NMD pathway, inhibiting further translation and recruiting other RNA-degrading enzymes.^45^ UPF1 has also been implicated in serving an antiviral role in the context of alphaviral infection.^46^ The authors of this study found that depletion of UPF1 or two other NMD components promotes viral replication, and specifically depleting UPF1 likely stabilizes incoming viral RNA genomes.^46^

We first validated that TRIM25 interacts with UPF1. To do so, we transfected V5-tagged UPF1 into TRIM25 inducible cell lines, then induced for TRIM25-WT or -R54P expression with dox, and performed a FLAG IP to pull down TRIM25. We found that UPF1 is robustly detected only when TRIM25 is induced (Fig. 4A), validating the TRIM25-UPF1 interaction identified in our co-IP/MS. To test the hypothesis that TRIM25 ubiquitinates UPF1, we co-transfected V5-tagged UPF1 with HA-Ub into our TRIM25 inducible cell lines and induced TRIM25 expression. We then performed a V5 IP and probed for the presence of ubiquitinated UPF1. We found that UPF1 is more robustly mono-ubiquitinated only in the presence of TRIM25-WT and not ligase-deficient TRIM25-R54P (~50% more by ImageJ quantification, Fig. 4B), suggesting that TRIM25 mono-ubiquitinates UPF1. We then identified putative ubiquitination sites by selecting residues that are both identified in a previously published ubiquitinome^47^ and predicted via UbPred to be ubiquitinated (Score > 0.70),^48^ and mutated these sites to arginine (K281R, K592R). Whereas ubiquitination is unchanged in UPF1 K281R, the introduction of K592R abolishes UPF1 ubiquitination in the presence of TRIM25-WT (Fig. 4C). Together, these results validate our co-IP/MS identification of UPF1 as a novel TRIM25 substrate.

**Figure 4.**
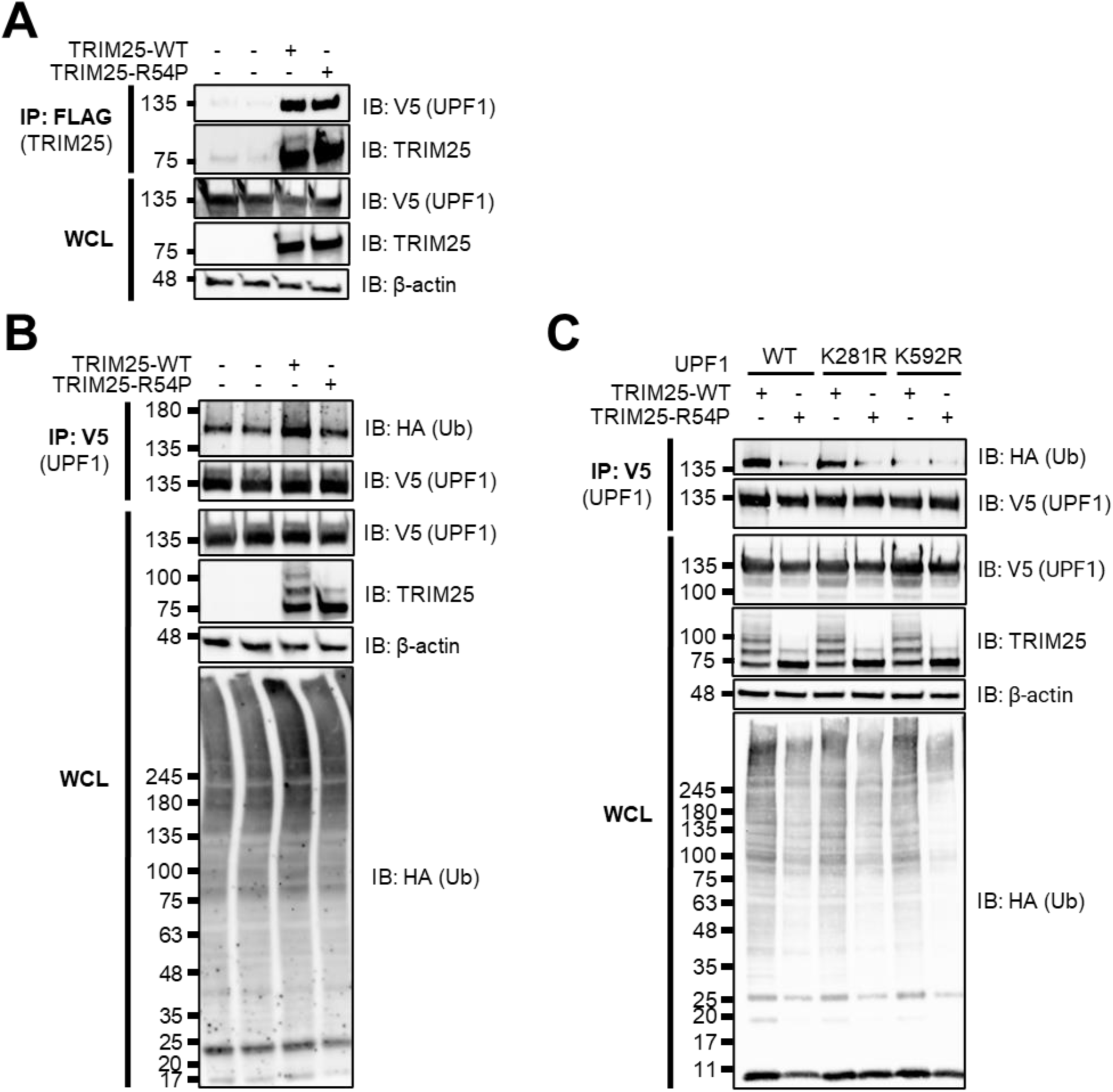
TRIM25 interacts with and mono-ubiquitinates UPF1. **(A)** Western blot of TRIM25 inducible cells transfected with V5-tagged UPF1 in the presence or absence of 1 μg/mL dox. Lysates were subjected to FLAG IP. Data are representative of three independent experiments. **(B-C)** Western blot of TRIM25 inducible cells transfected with **(B)** V5-UPF1 or **(C)** V5-UPF1 mutants (K281R, K592R) and HA-Ub in the presence of 1 μg/mL dox. Lysates were subjected to V5 IP. Data are representative of three independent experiments.

### TRIM25 polyubiquitinates NME1 but only interacts with endogenous, not ectopically expressed NME1

In addition, we identified NME1 as one of the most enriched TRIM25-R54P interactors in the presence of SINV infection (Table 3, log2FoldChange 3.4 – 4.8). NME1 is a nucleoside diphosphate kinase and a major synthesizer of non-ATP nucleoside triphosphates, perhaps best characterized in its role in inhibiting cell migration and proliferation of tumor cells via inhibition of MAPK signaling.^49^ However, the role of NME1 in viral replication is not well studied.^50^

Given its well-characterized role as a metastatic suppressor, we decided to validate NME1 as a TRIM25 ubiquitination substrate. We first set out to validate TRIM25 interaction with NME1 as identified in our co-IP/MS (Tables 1-3). To do so, we transfected myc-tagged NME1 or UPF1 to serve as a positive control in our TRIM25 inducible lines, induced for TRIM25-WT or -R54P expression, and performed a FLAG IP to pull down TRIM25. We then probed for any associated UPF1 or NME1. While we saw robust association of UPF1 with both TRIM25-WT and -R54P in line with our previous results (Fig. 4A, 5A, MW ~135 kDa), we did not identify NME1 (Fig. 5A, MW 20-25 kDa). We also performed the reverse IP where we pulled down myc-tagged NME1, but were unable to find any TRIM25 interacting with NME1 (Fig. 5B). We hypothesized that this lack of TRIM25-NME1 interaction could be due to functional differences between ectopically expressed myc-NME1 and endogenous NME1, given our successful validation of the other robust TRIM25 interactors from our co-IP/MS, G3BP and UPF1 (Fig. 3-4). To test this hypothesis, we performed a FLAG IP using our TRIM25 inducible lines and probed for co-IP of endogenous NME1 along with endogenous G3BP and UPF1 as positive controls. In line with our co-IP/MS results, endogenous G3BP, UPF1, and NME1 enrich robustly with TRIM25 pulldown, despite a low level of non-specific binding of NME1 to the FLAG IP in TRIM25 KO 293T cells (Fig. 5C).

**Figure 5.**
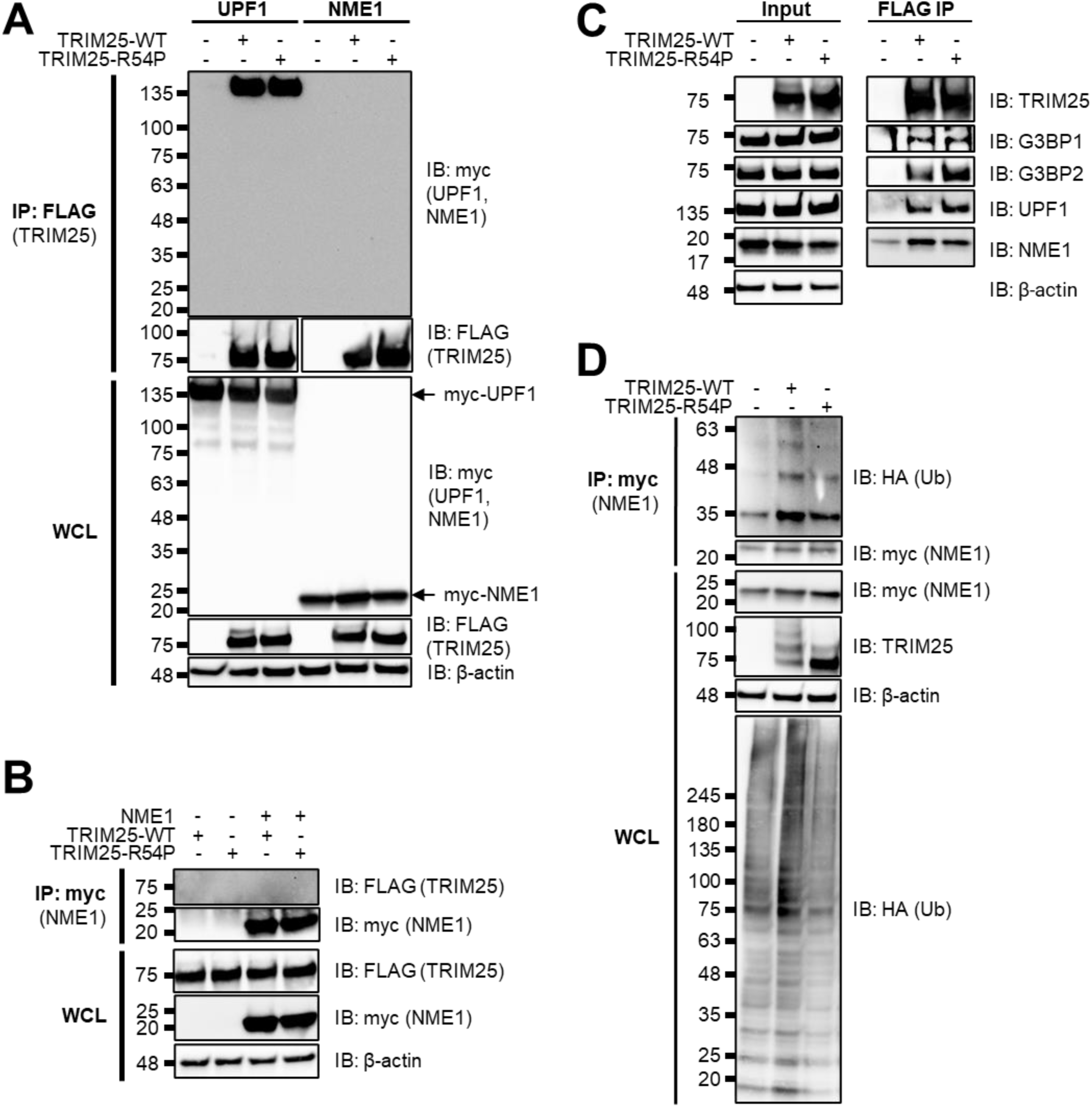
TRIM25 interacts with and polyubiquitinates NME1. **(A)** Western blot of TRIM25 KO and TRIM25 inducible cells transfected with myc-tagged UPF1 or NME1 in the presence of 1 μg/mL dox. Lysates were subjected to a FLAG IP. Data are representative of two independent experiments. **(B)** Western blot of TRIM25 inducible cells transfected with myc-NME1 in the presence or absence of 1 μg/mL dox. Lysates were subjected to a myc IP. Data are representative of two independent experiments. **(C)** Western blot of TRIM25 KO and TRIM25 inducible cells in the presence of 1 μg/mL dox. Lysates were subjected to a FLAG IP. Data are representative of two independent experiments. **(D)** Western blot of TRIM25 KO and TRIM25 inducible cells treated with 1 μg/mL dox and transfected with myc-NME1 and HA-Ub-WT. Lysates were subjected to myc IP. Data are representative of three independent experiments.

To test whether TRIM25 ubiquitinates NME1, we transfected myc-tagged NME1 into TRIM25-WT and -R54P inducible cells in the presence or absence of dox and performed a myc IP. We found that NME1 is more robustly polyubiquitinated in the presence of TRIM25-WT as compared to TRIM25-R54P, although we cannot yet rule out the possibility that TRIM25 might mono-ubiquitinate NME1 at multiple sites (Fig. 5D). Taken together, these results suggest that robust TRIM25 interactors, such as G3BP, UPF1, and NME1 identified in our TRIM25 co-IP/MS function as bona fide TRIM25 substrates.

### TRIM25 antiviral activity is dependent on its ligase activity

Given our identification of diverse host factors as TRIM25 substrates (Fig. 3-5), many of which function in translational and RNA processes (Fig. 2E) and several of which have known roles in alphavirus replication, we hypothesized that TRIM25 ligase activity is critical to orchestrating an antiviral response.

We used TRIM25 inducible cell lines in the KO background (Fig. 2A) to characterize the requirement of ligase activity in TRIM25-mediated viral inhibition. We found that TRIM25-WT, which retains ligase activity, represses SINV replication, whereas ligase mutant TRIM25-R54P does not (Fig. 6A). Overexpression of TRIM25-WT (Fig. 6A, solid light blue line) dramatically represses SINV replication by 7-15 fold at earlier timepoints (6-12 hours post infection (h.p.i.)) to 43-52 fold at later timepoints (24-40 h.p.i.) compared to TRIM25 KO 293T cell lines (Fig. 6A, dotted lines). Interestingly, some replicates fail to initiate infection in the presence of TRIM25-WT, causing seemingly large variability in viral replication. In contrast, overexpression of ligase-deficient TRIM25-R54P (Fig. 6A, solid dark blue line) restores SINV replication to levels even higher than the TRIM25 KO background (Fig. 6A, dotted lines). Overexpressed TRIM25-R54P may act in a dominant negative manner by binding to and sequestering ZAP, preventing ZAP from interacting with its other co-factors. Similarly, we found that overexpression of TRIM25-WT robustly represses virion production by approximately 36-250 fold at 24-40 h.p.i. (Fig. 6B, solid light blue line) and viral translation by 6 fold at 6 h.p.i. (Fig. 6C, light blue). In contrast, overexpression of TRIM25-R54P restores both virion production (Fig. 6B, solid dark blue line) and viral translation (Fig. 6C, dark blue) to comparable levels as the TRIM25 KO background. We then asked whether ligase-deficient TRIM25-R54P remains active against other alphaviruses. We tested other Old World (Ross River virus, RRV; o’nyong-nyong virus, ONNV) and New World (Venezuelan equine encephalitis virus, VEEV) alphaviruses. TRIM25-WT remains potently antiviral against all alphaviruses tested, while overexpression of TRIM25-R54P either has no effect on or restores viral replication to levels higher than the TRIM25 KO background (Fig. 6D). Taken together, these data clearly demonstrate that TRIM25-dependent ubiquitination is required for inhibition of alphavirus replication, specifically through a block in viral translation.

**Figure 6.**
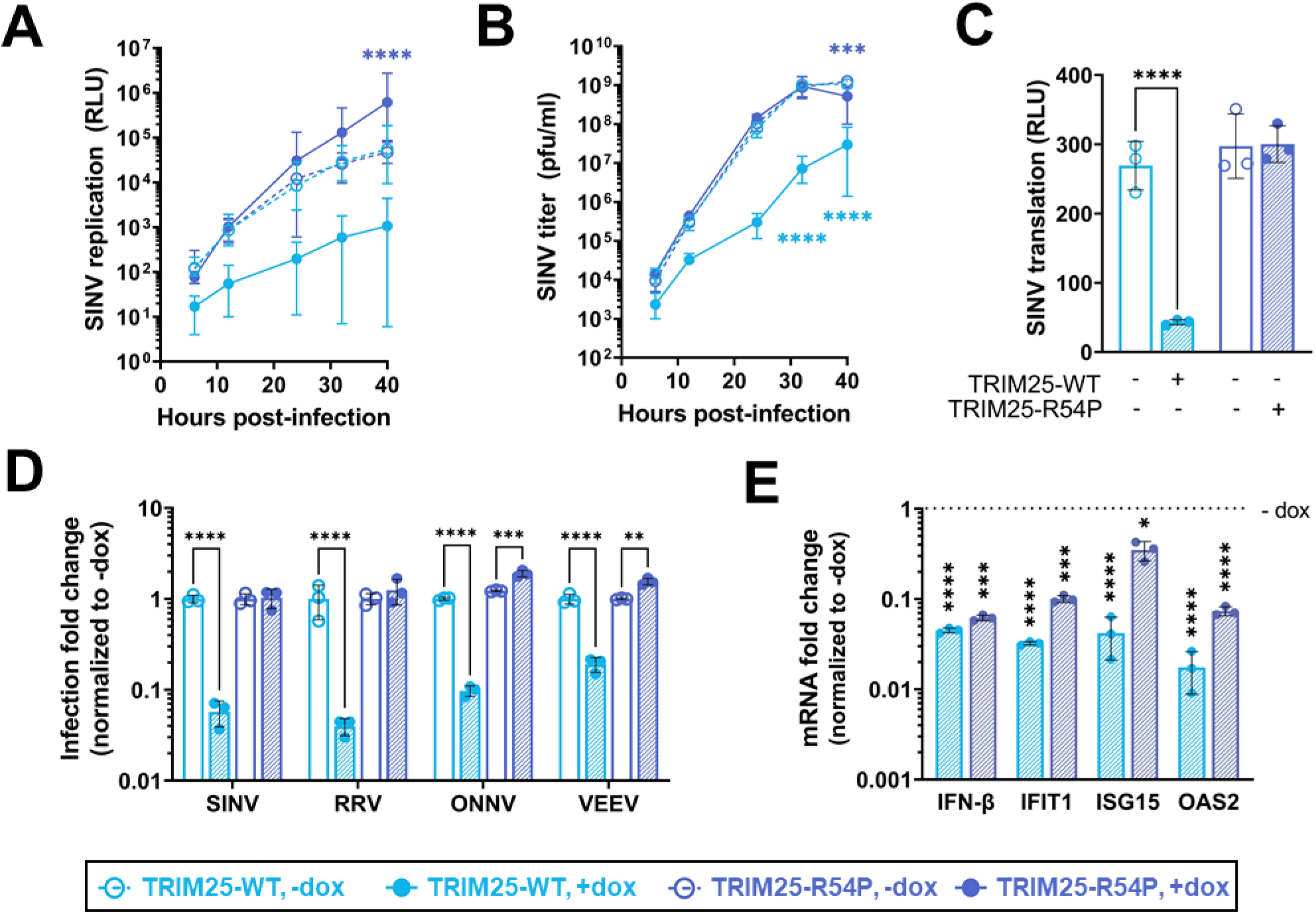
Point mutation in TRIM25 RING domain cripples TRIM25 antiviral activity. **(A-C)** Dox inducible TRIM25-WT or -R54P were integrated into TRIM25 KO 293T cells using the ePiggyBac (ePB) transposon, and induced for TRIM25-WT or -R54P expression at 1 μg/mL dox. Cells were infected with **(A)** SINV Toto1101/Luc at a MOI of 0.01 plaque forming unit (PFU)/cell, and lysed at 6, 12, 24, 32, and 40 hours post infection (h.p.i.); data combined from three independent experiments, error bars indicate range; or **(B**) Sindbis virus (SINV) Toto1101 at a MOI of 0.01 PFU/cell, harvesting supernatant at 6, 12, 24, 32, and 40 h.p.i. for plaque assays; data representative of two independent experiments, error bars indicate range; or **(C)** SINV Toto1101/Luc:ts6 at a MOI of 1 PFU/cell and lysed at 6 h.p.i. for measurement of luciferase activity; data representative of two independent experiments, error bars indicate standard deviation. **(D)** Percent infected cells (GFP+) at MOI of 0.01 PFU/cell (SINV 24 h.p.i.; Ross River virus (RRV) 24 h.p.i.; o’nyong-nyong virus (ONNV) 22 h.p.i.; Venezuelan equine encephalitis virus (VEEV) 10 h.p.i.) were normalized to that of the respective cell line without dox (set to one-fold). Asterisks indicate statistically significant differences, calculated using **(A-B, D)** Two-way ANOVA and Tukey’s multiple comparisons test: **, p<0.01; ***, p<0.001; ****, p<0.0001; (light blue compares WT +/− dox, dark blue compares R54P +/− dox) or **(C)** Two-way ANOVA and Sidak’s multiple comparisons test: ****, p<0.0001. **(E)** TRIM25 inducible cells were treated with poly(I:C) in the presence or absence of dox, and RNA harvested for RT-qPCR analysis. mRNA levels of IFN/ISGs in TRIM25-WT or R54P were normalized to that of the respective cell line without dox (set to one-fold). Data representative of two independent experiments. Asterisks indicate statistically significant differences as compared to the −dox condition (Two-way ANOVA and Sidak’s multiple comparisons test: *, p<0.05; ***, p<0.001; ****, p<0.0001).

### TRIM25-mediated viral inhibition is independent of changes in the type I IFN response

To exclude the complementary possibility that TRIM25 is exerting antiviral effects through affecting type I IFN or ISG production, we quantified the mRNA of IFN-β and the prominent ISGs IFIT1, ISG15, and OAS2 in the presence of poly(I:C), a dsRNA mimetic and stimulator of innate immune signaling. If TRIM25 antiviral activity is mediated through a strengthened IFN response, we would expect that both IFN and ISG production to increase when TRIM25-WT is induced and to be lower in the presence of TRIM25-R54P due to its defective antiviral activity. Poly(I:C) stimulation works well, inducing IFN-β and ISGs robustly in the absence of TRIM25 (>10 fold over absence of poly(I:C); data not shown). Surprisingly, we found that overexpression of either TRIM25-WT or TRIM25-R54P significantly suppresses production of IFN-β, IFIT1, ISG15, and OAS2 mRNA in the presence of poly(I:C) (Fig. 6E). We also observed that induction of TRIM25-WT results in a more drastic suppression of the ISGs as compared to TRIM25-R54P (Fig. 6E), which could be due to higher viral replication in the presence of TRIM25-R54P (Fig. 6A-B), leading to a higher type I IFN response in the TRIM25-R54P inducible cell line. Still, these data support our hypothesis that TRIM25 antiviral activity is not mediated through the IFN response.

### Identifying TRIM25-R54P specific interactors as critical for viral inhibition

As we showed that the loss of antiviral activity of TRIM25-R54P does not correlate with the levels of IFN and ISG expression, suggesting a direct consequence of TRIM25-mediated ubiquitination of target proteins, we then decided to examine TRIM25-R54P interactors identified in our co-IP/MS (Tables 1,3) that are not consistently present in the TRIM25-WT enrichment. These candidate proteins likely exhibit weaker or more transient interactions with TRIM25 and are ubiquitinated by TRIM25. We hypothesized that if any of these interactors are critical for TRIM25 antiviral activity, loss of their expression would result in increased viral replication even in the presence of overexpressed TRIM25-WT. While we initially also assessed a subset of ribosomal proteins identified as TRIM25-R54P interactors, their knockdown results in high cytotoxicity and therefore are excluded from subsequent analyses (data not shown). We validated most of the TRIM25-R54P interactors that are not present on the TRIM25-WT list (Tables 1-4) in the absence (Fig. 7A) or presence of viral infection (Fig. 7B).

**Figure 7.**
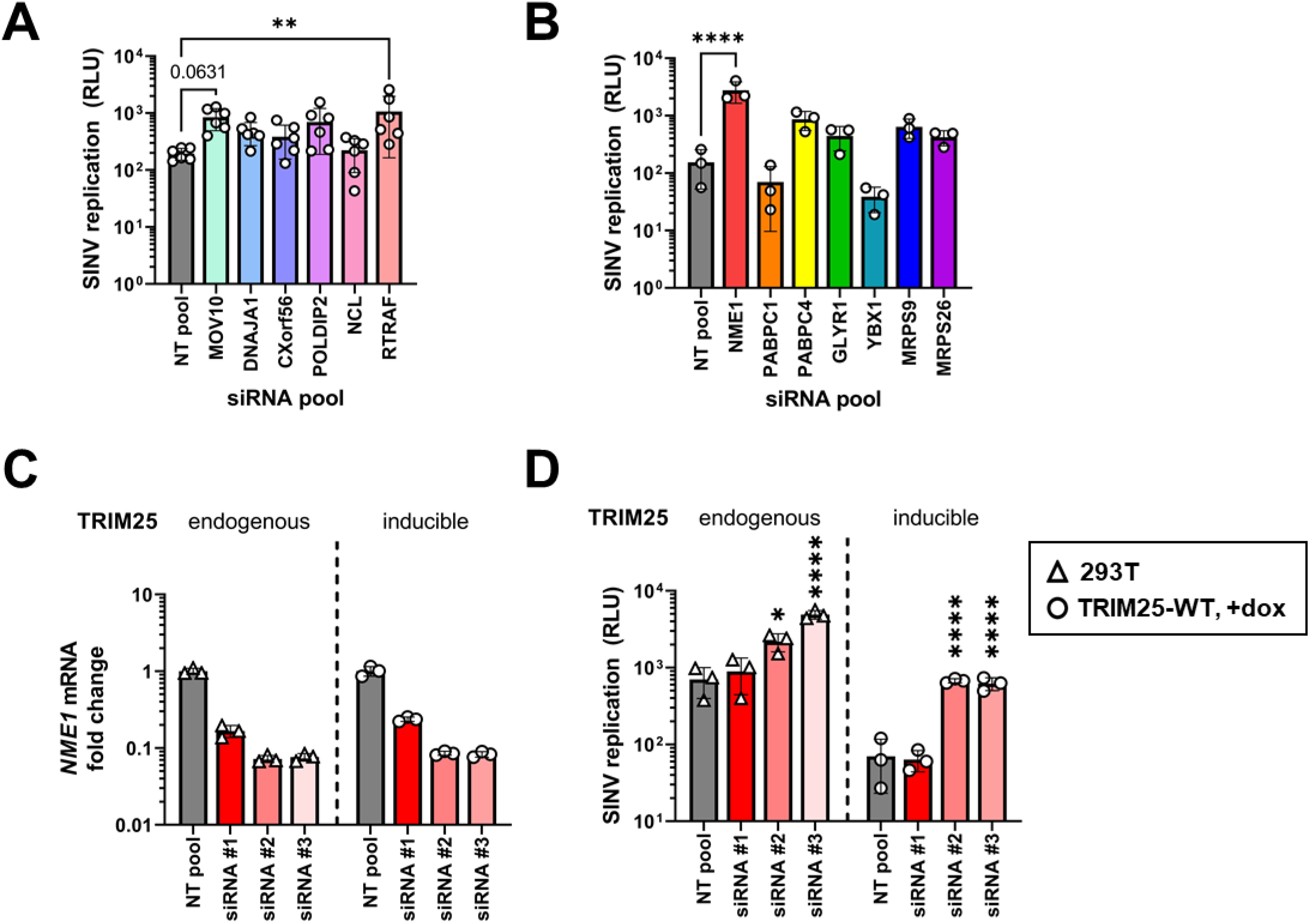
Knocking down TRIM25-R54P-specific interactors identifies essential substrates for TRIM25 antiviral activity. **(A-B)** TRIM25 inducible cells transfected with pooled siRNAs for TRIM25-R54P specific hits in the **(A)** absence or **(B)** presence of viral infection were induced for TRIM25-WT expression at 1 μg/mL dox), and infected with Toto1101/luc at MOI of 0.01 PFU/cell. Cells were lysed at 24 h.p.i. for measurement of luciferase activity. Asterisks indicate statistically significant differences as compared to the NT pool siRNA (One-way ANOVA, Dunnett’s multiple comparison test; **, p<0.01; ****, p<0.0001). Unlabeled comparisons are not significant. Data are either **(A)** pooled from or **(B)** representative of two independent experiments. **(C-D)** Parental 293T cells (TRIM25: endogenous) or TRIM25 inducible (TRIM25: inducible) cells were transfected with individual siRNAs for *NME1*, induced for TRIM25-WT expression at 1 μg/ml dox, and **(C)** had RNA extracted for RT-qPCR analysis or **(D)** infected with Toto1101/luc at MOI of 0.01 PFU/cell. Cells were lysed at 24 h.p.i. for measurement of luciferase activity. Asterisks indicate statistically significant differences as compared to the NT pool for each cell line. (One-way ANOVA, Dunnett’s multiple comparison test; *, p<0.05; ****, p<0.0001). Data are representative of two independent experiments.

While knockdown of multiple interactors trends towards restoring SINV replication, only loss of RTRAF (Table 1, log2FoldChange 1.6 – 1.8) and NME1 (Table 3, log2FoldChange 3.4 – 4.8) significantly restores SINV replication (Fig. 7A-B). Moreover, knockdown of MOV10 (Table 1, log2FoldChange 4.5 – 4.9) approaches significant restoration of SINV replication (Fig. 7A, p=0.0631).

Given that loss of NME1 results in the most significant restoration of SINV replication (Fig. 7B) and is ubiquitinated by TRIM25 (Fig. 5D), we de-convoluted its siRNA pool in both our inducible TRIM25-WT cell line and in the parental 293T cell line with endogenous TRIM25 and ZAP expression. There, we observed that the degree of *NME1* mRNA knockdown positively correlates with increase of viral replication (Fig. 7C-D), supporting a role for NME1 in TRIM25-dependent alphaviral inhibition. Altogether, these results suggest that the antiviral activity of TRIM25 is mediated by multiple substrates. Though knockdown of most individual interactors on their own does not significantly restore SINV replication, the fact that several have demonstrated a phenotype implies that together they may have a larger impact on viral replication. Further studies need to be performed to determine their synergistic effects on viral infection and functional consequences of their ubiquitination by TRIM25.

## DISCUSSION

Many TRIMs are involved in and ubiquitinate components of multiple cellular and antiviral processes.^3–6^ In this study, we set out to identify TRIM25 substrates by generating a point mutation in the TRIM25 RING domain, R54P, which is predicted to abolish its interaction with E2 carrier enzymes and is sufficient to cripple TRIM25 ligase activity (Fig. 1). We reported identification of TRIM25 substrates involved in nucleic acid metabolism and translation (Fig. 2E), in line with its role in blocking viral translation.^20^ We characterized the ubiquitination of the most enriched TRIM25-R54P interactors G3BP (Fig. 3), UPF1 (Fig. 4), and NME1 (Fig. 5), representing proteins with essential cellular functions, some of which with prior involvement in alphavirus infection.^46,51^ We also used the TRIM25-R54P mutant to definitively show the critical role of ubiquitination in TRIM25 antiviral activity that is independent of IFN production and signaling (Fig. 6). We then examined proteins that display a preference for association with TRIM25-R54P under mock and viral infection conditions, and found that several of these are necessary for TRIM25 antiviral activity (Fig. 7), identifying them as potential TRIM25 substrates mediating viral inhibition. Our results suggest that targeting of any single substrate by TRIM25 is insufficient to mediate the entirety of its cellular and antiviral activities, illustrating the powerful, multi-faceted role of this ubiquitination network in diverse biological processes.

We propose that the success of this “substrate trapping” approach in identifying TRIM25 ubiquitination substrates hinges on preservation of protein structure. Previous reports that unearthed the importance of TRIM25 ligase activity in the ZAP antiviral response depend on either deleting the entire TRIM25 RING catalytic domain or disrupting formation of the zinc finger motif, potentially having an adverse effect on protein folding overall and potentially affecting other TRIM25 cellular functions or interactions.^20,21^ The R54P point mutation we generated has been demonstrated to preserve protein structure and cognate interactions in other contexts,^29^ instilling greater credibility in our identification of novel TRIM25 substrates. Moreover, this mutation is predicted to abolish the E3 ligase-E2 conjugating enzyme interaction,^29^ preventing any downstream ubiquitination events and thus prolonging transient ligase-substrate interactions. The TRIM25-R54P specific hits may have weaker, more transient, or infection-specific interactions not easily detected by the conventional co-IP/MS approach. Other “substrate trapping” approaches depend on fusing a polyubiquitin binding domain to the ligase of interest,^52^ which may either disrupt native protein-protein interactions or result in false-positive identification of ubiquitinated proteins. Moreover, this type of approach would fail to identify substrates that are not polyubiquitinated, given that ligases can mono or multi-monoubiquitinate their substrates.^53^

For the first time, we identified G3BP1/2, UPF1, and NME1 as bona fide TRIM25 substrates (Fig. 3-5). Furthermore, we were able to characterize TRIM25 polyubiquitination of G3BP as primarily utilizing K63 linkages (Fig. 3C). This type of linkage is commonly used to build signaling scaffolds, as TRIM25 does to activate RIG-I, and could potentially play a role in either SG assembly or disassembly by recruiting SG components in the former or generating steric hindrance in the latter. Additionally, our validation of K592 as a monoubiquitination site on UPF1 (Fig. 4C) overlaps with a predicted acetylation site on the same residue, and neighbors a predicted phosphorylation site at T595, potentially modulating these other post-translational modifications of UPF1.^54^ These residues lie within the AAA ATPase domain of UPF1, suggesting that ubiquitination of UPF1 by TRIM25 might affect its ATP hydrolysis, thus hindering UPF1 in its NMD target discrimination and efficient translation termination.^55,56^ Interestingly enough, G3BP1 and UPF1 cooperate to mediate structure-mediated RNA decay.^57^ It is entirely possible that TRIM25-mediated ubiquitination could affect this process by modulating their interaction with one another, though further experiments are required to explore this hypothesis. Furthermore, NME1 has been previously demonstrated to be ubiquitinated and subsequently targeted for degradation by the E3 ligase SCF-FBXO24^58^. Seeing as TRIM25 is able to modify G3BP with both proteolytic K48- and non-proteolytic K63-polyubiquitin linkages, TRIM25 may also be targeting NME1 for degradation, thereby hindering nucleotide synthesis and general RNA metabolic processes.

We also utilized the TRIM25-R54P mutant to define the requirement for ligase activity in TRIM25 inhibition of alphavirus replication. We found that TRIM25 ligase activity is absolutely required for its inhibition of diverse alphaviruses through a block in viral translation. Interestingly, overexpression of both TRIM25-WT and -R54P results in a dampened IFN response in our hands (Fig. 6E), contrasting with the previously established role of TRIM25 in activating RIG-I signaling and implicating TRIM25 as a negative regulator of the type I IFN response.^59^ Moreover, TRIM25-R54P with a complete loss of antiviral activity actually exhibits relatively more production of IFN and a subset of ISG mRNAs (Fig. 6E), which may be indicative of higher viral replication overall (Fig. 6A-B). Still, these data together strongly suggest that the robust TRIM25 antiviral activity against alphaviruses is not mediated through an augmented IFN response, but through its ligase activity and subsequent ubiquitination network.

Our examination of the contribution of a subset of TRIM25-R54P specific interactors to TRIM25 antiviral activity has yielded several hits, namely RTRAF (Fig. 7A) and NME1 (Fig. 7B). RTRAF, also known as hCLE or C14orf166, is an RNA binding protein involved in cellular transcription, translation, and RNA transport, and is required for influenza virus replication.^60–62^ Notably, RTRAF is a member of a cap-binding complex that activates mRNA translation.^61^ Given RTRAF’s role in facilitating translation of mRNAs, it is therefore tempting to speculate that RTRAF may be required for translation of alphavirus RNA, and TRIM25-mediated ubiquitination of RTRAF may affect its ability to do so. The novel bona fide TRIM25 substrate NME1, which functions as a major synthesizer of non-ATP nucleoside triphosphates, upon ubiquitination may inhibit alphavirus replication via a similar mechanism as the potent restriction factor SAMHD1, which depletes deoxynucleotide pools, effectively preventing replication of varied DNA viruses and reverse transcription of HIV-1.^63^ On the other hand, TRIM25-mediated ubiquitination of NME1 may inhibit its metastatic suppressor activities, potentially serving as a novel mechanism for TRIM25’s previously described roles in carcinogenesis. Further studies need to be carried out to elucidate the functional consequences of these TRIM25 substrates in blocking viral translation and other cellular processes.

The novelty of this work lies within our innovative approach to uncover a multifaceted ubiquitination network involved in mediating TRIM25 cellular and antiviral functions. Many questions remain unanswered as to how TRIM25-mediated ubiquitination modulates the activity of these substrates. In contrast to the more binary consequences of K48-linked dependent degradation, other types of ubiquitin linkage may effect more nuanced cellular changes by modulating substrate activity and localization.^53^ Given TRIM25 proclivity for K63 linkages in the context of alphavirus infection and innate immunity,^19–21^ we are tempted to speculate that TRIM25 eschews a simple degradation approach in favor for a more nuanced modulation of substrate activity and localization. Current therapeutics that harness E3 ligases focus on their degradative power, generating compounds that bring ligases in close proximity to a target protein for degradation.^64^ Further research is warranted to explore the utility of alternate modes of ubiquitination in biological therapeutics.

## AUTHOR CONTRIBUTIONS

E.Y. and M.M.H.L. conceptualized and designed the study. E.Y. and S.H. performed experiments. Y.A. and J.W. performed co-IP/MS experiments and analyses. G.I. provided critical information and feedback concerning G3BP experiments. E.Y. wrote the first draft of the manuscript. M.M.H.L. provided critical feedback, and E.Y. and M.M.H.L. edited subsequent drafts. All authors contributed to manuscript revision, read, and approved the submitted version.

## ACKNOWLEDGMENTS

We thank Dr. Bill Schneider (Rockefeller University) for help with alignment of TRIM25 KO sequences and Drs. Douglas Black, Irvin Chen, and Oliver Fregoso (UCLA) for critical reading of the manuscript. We also thank the UCLA Proteome Research Center for their services. RT-qPCR and flow cytometry was performed in the UCLA AIDS Institute that is supported by the James B. Pendleton Charitable Trust and the McCarthy Family Foundation. Molecular graphics and analyses performed with UCSF Chimera, developed by the Resource for Biocomputing, Visualization, and Informatics at the University of California, San Francisco, with support from NIH P41-GM103311. This work was supported in part by National Institute of Health (NIH) grant (R01AI158704; M.M.H.L.), UC Cancer Research Coordinating Committee Faculty Seed Grant (CRN-20-637544; M.M.H.L.), UCLA AIDS Institute and Charity Treks 2019 Seed Grant (M.M.H.L.), Johanna and Joseph H. Shaper Family Chair (M.M.H.L.), Ruth L. Kirschstein Multidisciplinary Training Grant in Microbial Pathogenesis (NRSA AI007323; E.Y.)., Warsaw Fellowship (E.Y.), and Whitcome Fellowship (E.Y.).

## MATERIALS AND METHODS

Key resources table.

| Reagent type (species) or resource | Description | Source or reference | Identifiers | Additional information |
| --- | --- | --- | --- | --- |
| Antibody | anti-actin-HRP, mouse monoclonal | Sigma-Aldrich | A3854 | WB 1:20,000 |
| Antibody | anti-EFP/TRIM25, mouse monoclonal | BD Biosciences | 610570 | WB 1:5,000 |
| Antibody | anti-FLAG, mouse monoclonal | Sigma-Aldrich | F1804 | WB 1:20,000 |
| Antibody | anti-G3BP1, mouse monoclonal | Santa Cruz | sc-365338 | WB 1:500 |
| Antibody | anti-G3BP2, rabbit polyclonal | Assay Biotech | C18193 | WB 1:1,000 |
| Antibody | anti-HA, rat monoclonal | Roche Life Science | 3F10 | WB 1:1,000 |
| Antibody | anti-myc, rabbit polyclonal | Cell Signaling Technology | 2272S | WB 1:2,500 |
| Antibody | anti-NM23A (NME1), rabbit monoclonal | Abcam | ab171935 | WB 1:10,000 |
| Antibody | anti-UPF1, rabbit monoclonal | Cell Signaling Technology | 12040 | WB 1:1,000 |
| Antibody | anti-V5, mouse monoclonal | Invitrogen | MA5-1523 | WB 1:5,000 |
| Antibody | donkey anti-rat HRP | Jackson ImmunoResearch | 712-035-153 | WB 1:20,000 |
| Antibody | goat anti-mouse HRP | Jackson ImmunoResearch | 115-035-146 | WB 1:20,000 |
| Antibody | goat anti-rabbit HRP | Thermo Fisher Scientific | 31462 | WB 1:20,000 |
| Chemical compound, drug | poly(I:C) HMW | InvivoGen | tlrl-pic |  |
| Chemical compound, drug | Poly-L-lysine hydrobromide | Sigma-Aldrich | P2636 |  |
| Chemical compound, drug | Roche cOmplete Mini, EDTA-free Protease Inhibitor Cocktail | Sigma-Aldrich | 11836170001 |  |
| Commercial assay or kit | DharmaFECT 1 Transfection Reagent | Horizon Discovery | T-2001-01 |  |
| Commercial assay or kit | Dynabeads™ Protein A for Immunoprecipitation | Invitrogen | 10-002-D |  |
| Commercial assay or kit | EZview Red ANTI-FLAG M2 Affinity Gel | Sigma-Aldrich | F2426 |  |
| Commercial assay or kit | EZview Red ANTI-MYC M2 Affinity Gel | Sigma-Aldrich | E6654 |  |
| Commercial assay or kit | KOD Hot Start Master Mix | Sigma-Aldrich | 71842 |  |
| Commercial assay or kit | Luna Universal qPCR Master Mix | New England Biolabs | M3003X |  |
| Commercial assay or kit | Mini-PROTEAN TGX Gels, 4-15%, 15 well | Bio-Rad | 4568086 |  |
| Commercial assay or kit | NuPAGE MOPS SDS running buffer | Invitrogen | NP0001 |  |
| Commercial assay or kit | ProSignal Full-Range Prestained Protein Ladder | Genesee Scientific | 83-650 |  |
| Commercial assay or kit | ProSignal Pico ECL Reagent | Genesee Scientific | 20-300B |  |
| Commercial assay or kit | Protoscript II First Strand cDNA Synthesis Kit | New England Biolabs | E6560L |  |
| Commercial assay or kit | Q5 Site-Directed Mutagenesis Kit | New England Biolabs | E0552S |  |
| Commercial assay or kit | Quick-DNA Miniprep-Plus kit | Zymo Research | D4068 |  |
| Commercial assay or kit | Quick-RNA kit | Zymo Research | R1055 |  |
| Commercial assay or kit | QuikChange II XL Site-Directed Mutagenesis Kit | Agilent | 210518 |  |
| Commercial assay or kit | RNeasy mini kit | Qiagen | 74104 |  |
| Commercial assay or kit | XtremeGENE9 Transfection Reagent | Sigma-Aldrich | 6365787001 |  |
| Sequence-based reagent | DNA primers for molecular cloning | This work | See Supplementary file 1 |  |
| Sequence-based reagent | RT-qPCR oligonucleotides | PrimerBank | See Supplementary file 3 | https://pga.mgh.harvard.edu/primerbank/ |
| Sequence-based reagent | siRNA | Ambion | See Supplementary file 2 |  |
| Software, algorithm | Database for Annotation, Visualization and Integrated Discovery v6.8 | Frederick National Laboratory for Cancer Research, Frederick, MD |  | https://david.ncifcrf.gov/home.jsp |
| Software, algorithm | EnhancedVolcano | Clinical Bioinformatics Research LTD, United Kingdom |  | https://github.com/kevinblighe/EnhancedVolcano |
| Software, algorithm | FlowJo | BD Biosciences, Franklin, NJ |  | https://flowjo.com |
| Software, algorithm | Geneious Prime (2021.2) | Biomatters, San Diego, CA |  | https://geneious.com |
| Software, algorithm | GraphPad Prism 9 (v.9.2.0) | GraphPad Software, San Diego, CA |  | https://graphpad.com |
| Software, algorithm | ImageJ | National Institutes of Health, Bethesda, MD |  | https://imagej.net/ |
| Software, algorithm | RStudio software (v.1.4.1106) | RStudio, Boston, MA |  | https://rstudio.com |
| Software, algorithm | UCSF Chimera | University of California, San Francisco, San Francisco, CA |  | https://www.rbvi.ucsf.edu/chimera/ |

### Cell culture, viruses, and infections

ZAP KO 293T cells (clone 89) and its respective parental 293T cells were generously provided by Dr. Akinori Takaoka at Hokkaido University^65^. 293T (parental, ZAP KO, and TRIM25 KO (see below) with or without inducible expression of TRIM25) were cultured in Dulbecco’s Modified Eagle Medium (DMEM; Thermo Fisher Scientific, Waltham, MA) supplemented with 10% fetal bovine serum (FBS; Avantor Seradigm, Radnor, PA). Baby hamster kidney 21 (BHK-21; American Type Culture Collection, Manasass, VA) cells were cultured in Minimal Essential Media (Thermo Fisher Scientific) supplemented with 7.5% FBS.

Wild-type SINV (Toto1101), temperature-sensitive SINV (Toto1101/Luc:ts6), SINV expressing firefly luciferase (Toto1101/Luc), SINV expressing EGFP (TE/5’2J/GFP), CHIKV vaccine strain 181/clone 25 (generously provided by Scott Weaver, The University of Texas Medical Branch at Galveston), ONNV expressing EGFP (generously provided by Dr. Steve Higgs, Kansas State University), RRV expressing EGFP (generously provided by Dr. Mark Heise, University of North Carolina), and VEEV vaccine strain TC-83 (generously provided by Dr. Ilya Frolov, University of Alabama at Birmingham) have been previously described.^66–71^ Viral stocks and titers for multiplicity of infection (MOI) calculations were generated in BHK-21 cells as previously described.^66^ Viral infections and plaque assays were performed as previously described.^66^ TRIM25 inducible cells (see below) were induced for TRIM25 expression and infected with EGFP expressing viruses at a MOI of 0.01, harvested at 10-24 h.p.i., and fixed in 1% paraformaldehyde for flow cytometry analysis. Data was acquired using a MACSQuant Analyzer 10 (Miltenyi Biotec, Auburn, CA) and analyzed using FlowJo (BD Biosciences, Franklin Lakes, NJ). Percent infected (GFP+) cells was calculated and normalized to the −dox condition of each respective cell line.

### Plasmids and transfections

See Supplementary file 1 corresponding to Supplementary Table 1: Cloning primers. Addgene plasmids for HA-tagged ubiquitin (pRK5-HA-Ubiquitin-WT, #17608; pRK5-HA-Ubiquitin-K48, #17605; pRK5-HA-Ubiquitin-K63, #17606) and UPF1 (pCW57.1-Tet-UPF1WT, #99146) were used.^44,72^ Full-length TRIM25 was generously provided by Dr. Jae U. Jung at the University of Southern California.^19^ Dr. Gerald McInerney at the Karolinska Institutet, Sweden, generously provided pGFP-G3BP1 and pGFP-G3BP2a.^51^ The coding sequence of NME1 isoform a (NM_198175.1) was synthesized as a gene fragment (Integrated DNA Technologies, Coralville, IA), where the ends were flanked by restriction enzyme sites NotI and XbaI, and random nucleotides were incorporated to maintain the open reading frame. Dr. Oliver Fregoso kindly gifted us a pcDNA3.1-3XFLAG plasmid. The 3XFLAG tag was swapped out for a V5 tag or a myc tag using BamHI and HindIII restriction sites to generate V5-pcDNA3.1 or myc-pcDNA3.1, respectively. The pcDNA3.1-3XFLAG, pcDNA3.1-V5, and pcDNA3.1-myc vectors were used as an expression vector for TRIM25, UPF1, G3BP, and NME1. TRIM25 was cloned into pcDNA3.1-3XFLAG using XhoI and XbaI restriction sites, while UPF1, G3BP, and NME1 were cloned into either pcDNA3.1-V5 (UPF1) or pcDNA3.1-myc (G3BP and NME1) using the NotI and XbaI restriction sites. TRIM25 RING domain mutants (I15K, R54P, I15K/R54P) were generated by mutagenesis of pcDNA-3XFLAG-TRIM25 using the QuikChange II XL Site-Directed Mutagenesis Kit (Agilent Technologies, Santa Clara, CA), while the TRIM25-PTAA mutant was generated using the Q5 Site-Directed Mutagenesis Kit (New England Biolabs, Ipswich, MA), by performing sequential mutagenesis reactions to individually mutate each residue to alanine. TRIM25 was cloned into a 3XFLAG expressing ePiggyBac transposon plasmid at the ClaI and NotI restriction sites. All plasmids were verified by sequencing (Genewiz, South Plainfield, NJ).

Cells were transfected using XtremeGENE9 DNA Transfection Reagent (Roche Life Science, Basel, Switzerland) at a ratio of 3 μl to1 μg DNA according to the manufacturer’s instructions. Empty vectors (pcDNA3.1-3XFLAG, V5, or myc) were transfected as necessary to keep total plasmid amount in co-transfections constant.

### *TRIM25* targeting by CRISPR

The MIT Optimized CRISPR Design portal (<u>crispr.mit.edu</u>) and CHOPCHOP^73^ (chopchop.cbu.uib.no) were used to design guide RNAs (gRNAs) targeting exon 1 of the human *TRIM25* gene (Fig. S1A). The guide with the highest ranking in both scoring programs (5’-CGGCGCAACAGGTCGCGAACGGG-3’) was selected for cloning into the PX459 vector (Addgene, #62988), a non-lentiviral construct that also delivers Cas9.^74^ Oligos containing the gRNA sequences (5’-CACCGCGGCGCAACAGGTCGCGAAC-3’ and 5’-AAACGTTCGCGACCTGTTGCGCCGC-3’) were ligated and cloned into PX459 linearized with BbsI. 293T cells were transiently transfected with PX459 expressing TRIM25 gRNA and selected with 1 μg/mL puromycin the next day to eliminate untransfected cells. Following two days of puromycin selection, surviving cells were counted, diluted to 0.3 cell/well in a 96-well plate, and seeded in 10% FBS DMEM. Single cell clones were expanded and treated with or without puromycin. Clones sensitive to puromycin, indicating failure to integrate gRNA expressing vector, were harvested for immunoblot analysis to assess TRIM25 expression. Five clones (3, 6, 8, 9, and 10) were selected based on western blotting results indicating complete loss of TRIM25 protein expression (Fig. S1B). Viral replication within these clones was characterized by infection with a luciferase-expressing SINV (Toto1101/Luc). Clone #8 was selected for generation of TRIM25 inducible cell lines based on its intermediate viral replication phenotype (Fig. S1C), similar to previous TRIM25 siRNA data.^20^ A 600-bp amplicon flanking the gRNA targeting site was amplified from genomic DNA isolated from each clonal population using a Quick-DNA Miniprep Plus kit (Zymo Research, Irvine, CA) and KOD Hot Start Master Mix (Millipore Sigma). Amplicons from clone #8 were sent to Massachusetts General Hospital Center for Computational and Integrative Biology DNA Core for Complete Amplicon Sequencing, confirming that CRISPR targeting results in deletions in exon 1 of *TRIM25*, leading to frameshift mutations and premature stop codons in both alleles (Fig. S1D).

### Generation of TRIM25 inducible cell lines

To reconstitute TRIM25 expression (WT and R54P) in our TRIM25 KO 293T cell line (clone #8; see above for details), we used the enhanced PiggyBac (ePB) transposable element system provided by the Brivanlou laboratory at the Rockefeller University, as previously described.^75,76^ TRIM25 KO 293T cells were transfected with 1:1 ePB transposon vector encoding TRIM25-WT or TRIM25-R54P and the transposase plasmid. Two days post-transfection, 1.5 μg/mL of puromycin was used to select a population of TRIM25 KO 293T cells inducible for TRIM25-WT, -R54P, or -PTAA, which were then expanded and treated with different amounts of dox (0.01, 0.1, 1, and 10 μg/mL) to confirm TRIM25 inducible expression by immunoblotting.

### Mass spectrometry (MS)

To identify TRIM25 substrates, three 15-cm dishes per condition were seeded with 7.5×10^6^ TRIM25 inducible or TRIM25 KO 293T cells each in the presence of 1 μg/mL dox. Two days later, cells were mock infected or infected with Toto1101 at an MOI of 1 plaque forming unit (PFU)/cell. Six hours post infection, cells were trypsinized, spun down, and lysed in 3 mL of FLAG IP buffer. Supernatant was transferred to a new 15 mL tube and supplemented with 5 mL of FLAG IP buffer before incubating with 80 μL of anti-FLAG beads for 45 min at 4°C, rotating. Immunoprecipitates were washed three times in FLAG IP buffer before elution with 130 μL of 8M urea in 100 mM Tris-HCl, pH 8, shaken for 10 min at 1200 rpm. Supernatant was carefully transferred to a new tube and proteins were precipitated by addition of 4 volumes of −20°C acetone and incubation at 4°C overnight. After centrifugation at 16,100g for 30 min at 4°C, pellets were washed with −20°C acetone and centrifuged again.

Dried pellets were processed at the UCLA Proteomics Core. Protein samples were reduced and alkylated using 5mM Tris (2-carboxyethyl) phosphine and 10mM iodoacetamide, respectively, and then proteolyzed by the sequential addition of trypsin and lys-C proteases at 37°C as described.^77^ Digested peptides were resuspended in 5% formic acid and fractionated online using a 25cm long, 75 μM inner diameter fused silica capillary packed in-house with bulk C18 reversed phase resin (length, 25 cm; inner diameter, 75 μM; particle size, 1.9 μm; pore size, 100 Å; Dr. Maisch GmbH).^78^ The 140-minute water-acetonitrile gradient was delivered using a Dionex Ultimate 3000 UHPLC system (Thermo Fisher Scientific) at a flow rate of 300 nL/min (Buffer A: water with 3% DMSO and 0.1% formic acid and Buffer B: acetonitrile with 3% DMSO and 0.1% formic acid). Fractionated peptides were ionized and analyzed by tandem mass spectrometry (MS/MS) Orbitrap Fusion Lumos mass spectrometer (Thermo Fisher Scientific). Label-free quantitation was performed using the MaxQuant software package.^79^ The EMBL Human reference proteome (UP000005640 9606) was utilized for all database searches. Statistical analysis of MaxQuant output data was performed with the artMS Bioconductor^80^ package which performs the relative quantification of protein abundance using the MSstats Bioconductor package (default parameters). Intensities were normalized across samples by median-centering the log2-transformed MS1 intensity distributions. The abundance of proteins missing from one condition but found in more than 2 biological replicates of the other condition for any given comparison were estimated by imputing intensity values from the lowest observed MS1-intensity across samples and p-values were randomly assigned to those between 0.05 and 0.01 for illustration purposes. Significant hits were defined as interactors that possessed a log2FoldChange of >1.5 and a −log10Pvalue > 1.3.

### TRIM25 autoubiquitination and co-immunoprecipitation (co-IP) assay

To assess TRIM25 autoubiquitination or co-immunoprecipitation with proteins of interest, transfected or untransfected cells in 6-well plates were collected and lysed by rotating for 30 min at 4°C in FLAG IP buffer (100 mM Tris-HCl 8.0, 150 mM NaCl, 5 mM EDTA, 1 mM DTT, 5% glycerol, 0.1% NP-40) supplemented with a complete protease inhibitor cocktail (Roche), before spinning down at 14000 rpm for 15 min at 4°C. Anti-FLAG beads (EZview Red ANTI-FLAG M2 Affinity Gel, Sigma-Aldrich, St. Louis, MO) or anti-myc beads (EZview Red ANTI-MYC M2 Affinity Gel, Sigma-Aldrich) were equilibrated by washing 3 times in FLAG IP buffer. Three hundred μL of whole cell lysate (WCL) were incubated with 30 μL of anti-FLAG beads for 45 minutes at 4°C, rotating. Immunoprecipitates were washed 3 times with the FLAG IP buffer. Bound proteins were eluted with SDS loading buffer and boiled for 5 minutes for immunoblot analysis.

### Ubiquitination IP assay

To assess TRIM25 ubiquitination of putative substrates, immunoprecipitation was performed essentially as previously described.^20^ Briefly, cells were collected and lysed in 0.5% SDS buffer supplemented with complete protease inhibitor cocktail. Three hundred μL of WCL were diluted into 1X TNA buffer (0.25% Triton, 50 mM Tris-HCl, pH 7.5; 200 mM NaCl, 1 mM EDTA) + 2 mg/mL BSA. WCL containing V5-tagged substrates were then incubated with 1 μg of anti-V5 antibody overnight at 4°C. The next morning, 40 μL Protein A Dynabeads (Invitrogen, Waltham, MA) were added and incubated for 2 h at 4°C. WCL containing myc-tagged substrates were incubated directly with anti-myc beads for 45 minutes at 4°C, rotating. Following incubation with beads, both myc-tagged and V5-tagged immunoprecipitates were washed 3 times with 1X TNA buffer + 2 mg/mL BSA. Myc-tagged NME1 underwent an additional two washes with 1X TNA buffer only. Bound proteins were eluted with SDS loading buffer and boiled for 5 minutes for immunoblot analysis.

### Immunoblot analysis

Proteins were resolved through SDS-PAGE using 4-15% precast Mini-PROTEAN TGX Gels (Bio-Rad) and NuPAGE MOPS SDS running buffer (Thermo Fisher Scientific) before transferring to a PVDF membrane (Bio-Rad). Immunodetection was achieved with 1:5000 anti-ZAP (Abcam, Cambridge, United Kingdom); 1:5000 anti-TRIM25 (BD Biosciences), 1:100 anti-RIG-I (Cell Signaling Technology, Danvers, MA), 1:1000 anti-HA (Roche Life Sciences), 1:5000 anti-V5 (Invitrogen), 1:2500 anti-myc (Cell Signaling Technology), 1:20,000 anti-FLAG (Sigma-Aldrich), 1:500 anti-G3BP1 (Santa Cruz, Dallas, TX), 1:1000 anti-G3BP2 (Assay Biotech, Fremont, CA), 1:1000 anti-UPF1 (Cell Signaling Technology), 1:10,000 anti-NME1 (Abcam), and 1:20,000 anti-actin-HRP (Sigma-Aldrich). Primary antibodies were detected with 1:20,000 goat anti-mouse HRP (Jackson ImmunoResearch, West Grove, PA), 1:20,000 goat anti-rabbit HRP (Thermo Fisher Scientific), or 1:20,000 donkey anti-rat HRP (Jackson ImmunoResearch). Proteins were resolved on a 4-15% Mini-PROTEAN TGX gel (Bio-Rad, Hercules, CA) and visualized using ProSignal Pico ECL Reagent (Genesee Scientific, San Diego, CA) on a ChemiDoc (Bio-Rad). Quantification of western blots was performed using ImageJ.^81^

### siRNA knockdown

See Supplementary file 2 corresponding to Supplementary Table 2: siRNAs. Ambion Silencer siRNAs and nontargeting controls (Thermo Fisher Scientific) were reverse transfected with DharmaFECT 1 Transfection Reagent (Horizon Discovery, Cambridge, United Kingdom) according to manufacturer protocols. Briefly, siRNAs were mixed with DharmaFECT 1 Transfection Reagent (1:100 dilution in HBSS) and 50 μL of siRNA mix were added to each well in a 24 well plate, or 100 μL in a 12 well plate. 1.2 x 10^5^ cells were added per well in 250 μL in a 24 well plate or 2.4 x 10^6^ in 500 μL in a 12 well plate, for a final concentration of 25 nM siRNA and total volume of 300 μL. Plates that would be subjected to SINV infection were first poly-L-lysine treated. Cells were induced for TRIM25 expression using a final concentration of 1 μg/mL dox one day post-transfection, as applicable. Cells were harvested for RNA extraction for RT-qPCR to quantify gene knockdown (see below) or subjected to SINV infection 48 h post-transfection. To assess ISG induction in TRIM25 inducible cells upon poly(I:C) treatment, one day post-transfection cells were treated with 1 μg poly(I:C) HMW (InvivoGen, San Diego, CA) and 1 μg/mL dox per well. RT-qPCR data were normalized to the −dox condition for respective cell lines.

### Quantitative reverse transcription PCR (RT-qPCR)

See Supplementary file 3 corresponding to Supplementary Table 3: RT-qPCR primers. Total RNA was isolated from siRNA-treated cells using the RNeasy mini kit (Qiagen, Hilden, Germany) or the Quick-RNA kit (Zymo Research). 400 ng to 1 μg of input RNA was used as a template for reverse transcription using Protoscript II First Strand cDNA Synthesis Kit (NEB) and random hexamers, following manufacturer instructions. RT-qPCR was performed using 5 μL of 4 to 10-fold-diluted cDNA, and Luna Universal qPCR Master Mix (NEB) in the CFX Real-Time PCR system (Bio-Rad), courtesy of the UCLA Virology Core. qPCR conditions were as follows: initial denaturation step at 95 °C for 1 min, then 40 cycles of 95°C for 15 sec followed by 60°C for 30 sec, concluding with a final 10 sec at 60°C. A melt curve was then calculated by heating to 95°C incrementally by 0.5°C/s for 10 sec at each temperature. Transcript levels of ISGs and R54P specific interactors were determined by normalizing the target transcript CT value to the CT value of the RPS11 transcript, an endogenous housekeeping gene. Fold change was calculated using this normalized value relative to the average of cells treated with the NT siRNA control (CT method).

